# The capsule regulatory network of *Klebsiella pneumoniae* defined by density-TraDISort

**DOI:** 10.1101/404400

**Authors:** Matthew J. Dorman, Theresa Feltwell, David A. Goulding, Julian Parkhill, Francesca L. Short

## Abstract

*Klebsiella pneumoniae* infections affect infants and the immunocompromised, and the recent emergence of hypervirulent and multi-drug resistant *K. pneumoniae* lineages is a critical healthcare concern. Hypervirulence in *K. pneumoniae* is mediated by several factors, including the overproduction of extracellular capsule. However, the full details of how *K. pneumoniae* capsule biosynthesis is achieved or regulated are not known. We have developed a robust and sensitive procedure to identify genes influencing capsule production, density-TraDISort, which combines density gradient centrifugation with transposon-insertion sequencing. We have used this method to explore capsule regulation in two clinically-relevant *Klebsiella* strains; *K. pneumoniae* NTUH-K2044 (capsule type K1), and *K. pneumoniae* ATCC43816 (capsule type K2). We identified multiple genes required for full capsule production in *K. pneumoniae*, as well as putative suppressors of capsule in NTUH-K2044, and have validated the results of our screen with targeted knockout mutants. Further investigation of several of the *K. pneumoniae* capsule regulators identified – ArgR, MprA/KvrB, SlyA/KvrA and the Sap ABC transporter – revealed effects on capsule amount and architecture, serum resistance and virulence. We show that capsule production in *K. pneumoniae* is at the centre of a complex regulatory network involving multiple global regulators and environmental cues, and that the majority of capsule regulatory genes are located in the core genome. Overall our findings expand our understanding of how capsule is regulated in this medically-important pathogen, and provide a technology that can be easily implemented to study capsule regulation in other bacterial species.

**Importance:** Capsule production is essential for *K. pneumoniae* to cause infections, but its regulation and mechanism of synthesis are not fully understood in this organism. We have developed and applied a new method for genome-wide identification of capsule regulators. Using this method, many genes that positively or negatively affect capsule production in *K. pneumoniae* were identified, and we use these data to propose an integrated model for capsule regulation in this species. Several of the genes and biological processes identified have not previously been linked to capsule synthesis. We also show that the methods presented here can be applied to other species of capsulated bacteria, providing the opportunity to explore and compare capsule regulatory networks in other bacterial strains and species.

## Introduction

*Klebsiella pneumoniae* is an ubiquitous Gram-negative bacterium, found both in the environment and as an asymptomatic coloniser of the mucosal surfaces of mammals (1). *K. pneumoniae* is also an opportunistic pathogen, and can express a multitude of virulence factors which enable it to cause infections in humans (1–4). Historically associated with infections in the immunocompromised and in neonates (1, 5), focus has been directed to *K. pneumoniae* following the emergence of antimicrobial-resistant and hypervirulent lineages (6, 7). Hypervirulent lineages are a particular concern in a clinical setting, because they have the potential to cause infection in immunocompetent hosts (6, 8, 9), to metastasise (10), and to cause infections in unusual infection sites (11).

Numerous factors contribute to *K. pneumoniae* virulence, such as the ability to produce siderophores, fimbriae, lipopolysaccharide, and extracellular polysaccharide capsule (6, 12–14). Hypervirulence is associated with these factors, particularly with the overproduction of capsular polysaccharide (6, 12, 15, 16), and in the absence of these virulence factors, *K. pneumoniae* virulence is reduced or abolished (14, 17, 18). The ∼200 kilobase *K. pneumoniae* virulence plasmid, which is also associated with the hypervirulent phenotype (6), encodes siderophores such as aerobactin and salmochelin, and positive regulators of capsule biosynthesis (6, 19–21). More than 100 capsule locus types have been identified in *K. pneumoniae* (17), though the majority of clinical cases of *K. pneumoniae* infection are caused by strains of capsule types K1 and K2 (12, 22).

Excessive capsule production is strongly associated with hypervirulence in *K. pneumoniae* (23–25) and several studies have sought to identify genetic determinants of hypervirulence. For example, the mucoviscosity-associated gene *magA* (now named *wzy_K1* (26, 27)) was originally identified by transposon mutagenesis screening (28). The *rmpA* and *rmpA2* genes also affect hypermucoviscosity, and encode transcription factors that positively regulate the *K. pneumoniae* capsule biosynthesis locus (6, 15, 16, 29). These regulators can be either chromosomally-encoded or plasmid-borne. Although *rmpA* is correlated with hypervirulence, and strains lacking *rmpA* and aerobactin are avirulent in mice (12, 30), it has been shown that this increased virulence is a consequence of the hypermucoviscous phenotype conferred by *rmpA*, rather than the gene itself (24).

Capsule is also a potential therapeutic target in *K. pneumoniae*. Capsule-targeting monoclonal antibodies increased the killing of *K. pneumoniae* ST258 (an outbreak lineage) by human serum and neutrophils (31, 32) and limited the spread of a respiratory *K. pneumoniae* ST258 infection in mice. Specific capsule-targeting bacteriophage have been shown to clear or limit infections caused by *K. pneumoniae* strains of capsule types K1, K5 and K64 (33–35), and in some cases protection could also be achieved by treatment with the capsule depolymerase enzymes produced by these phage, rather than the phage itself. Translating these early-stage findings to the clinic is a priority as extensively drug-resistant *K. pneumoniae* strains become more prevalent (36).

Despite the absolute requirement for capsule in *Klebsiella* infections, its promise as a therapeutic target, and the connection between capsule overproduction and hypervirulence, the biosynthetic and regulatory mechanisms governing *K. pneumoniae* capsule have not yet been fully explored. Genetic screens and reverse genetics (28, 37–41), including transposon mutagenesis approaches (38), have been used to identify biosynthetic genes and activators of capsule production (including the *rmpA* and *magA* genes), and some of the cues that elevate capsule expression above basal levels have also been described – these include temperature, iron availability and the presence of certain carbon sources (12, 42). Defining the capsule regulatory network of *Klebsiella pneumoniae* in more detail would not only deepen our understanding of this pathogen’s interaction with the host environment, but could also inform efforts to target this factor with new therapeutics.

Here we employed transposon directed insertion sequencing (TraDIS) (43, 44) to identify genes influencing *K. pneumoniae* capsule production. We performed density-based selection on mutant libraries of both K1 and K2 capsule type *K. pneumoniae*, using a discontinuous density gradient (45). This approach allowed for the simultaneous selection of both capsulated and non-capsulated mutants from mutant libraries, without requiring growth of bacteria under a selective pressure. We have identified 78 genes which, when mutagenised by transposon insertion, reduce the ability of one or both of these strains to manufacture capsule. These included multiple genes not previously associated with capsule production in *K. pneumoniae*. We have also identified 26 candidate genes in NTUH-K2044 which cause an increase in capsulation when disrupted. Our results allow us to present an integrated model for capsule regulation in *K. pneumoniae*, and establish a technology for study of capsule production that is applicable to other bacterial species.

## Results

### Density gradient centrifugation separates bacterial populations based on capsule phenotype

*K. pneumoniae* capsule influences the centrifugation period required to pellet cells, and this property has been used to semi-quantitatively compare capsule production between different strains (38). Based on this observation, we speculated that *Klebsiella* cells of different capsulation states could be separated by density-based centrifugation, and that this method could be combined with TraDIS to screen for capsule-regulating genes. Discontinuous Percoll density gradients are routinely used to purify macrophages from complex samples (46), and have also been used to examine capsulation in *Bacteroides fragilis* and *Porphyromonas gingivalis* (45, 47). Tests with *K. pneumoniae* NTUH-K2044 (capsule type K1, hypermucoid, Fig. 1A), *K. pneumoniae* ATCC43816 (K2, hypermucoid) and a non-capsulated *E. coli* control showed that these strains differed in density and migrated to above 15%, above 35% and below 50% Percoll, respectively (Fig. S1A). Growth of the *Klebsiella* strains at 25 °C (which reduces capsule production) decreased their density, with NTUH-K2044 localising to above 35% Percoll, and ATCC43816 showing limited migration into the 35% layer. A third *K. pneumoniae* strain, RH201207 (non-hypermucoid, capsule type K106), was retained above 50% Percoll (Fig. S1A). We also examined the proportion of *Klebsiella* bacteria migrating to different locations within the density gradient (Fig. S1B). While the majority of *K. pneumoniae* NTUH-2044 bacteria were located above the 15% layer, a small fraction (4% viable count of the top) migrated to below the interface of this layer, suggesting some heterogeneity in capsule production. In both NTUH-K2044 and ATCC43816, only a very small number of cells were present below the 50% layer (Fig. S1B); these numbers could be partially due to cross-contamination during extraction of this part of the gradient, which was recovered by pipetting from the top.

**Figure 1:**
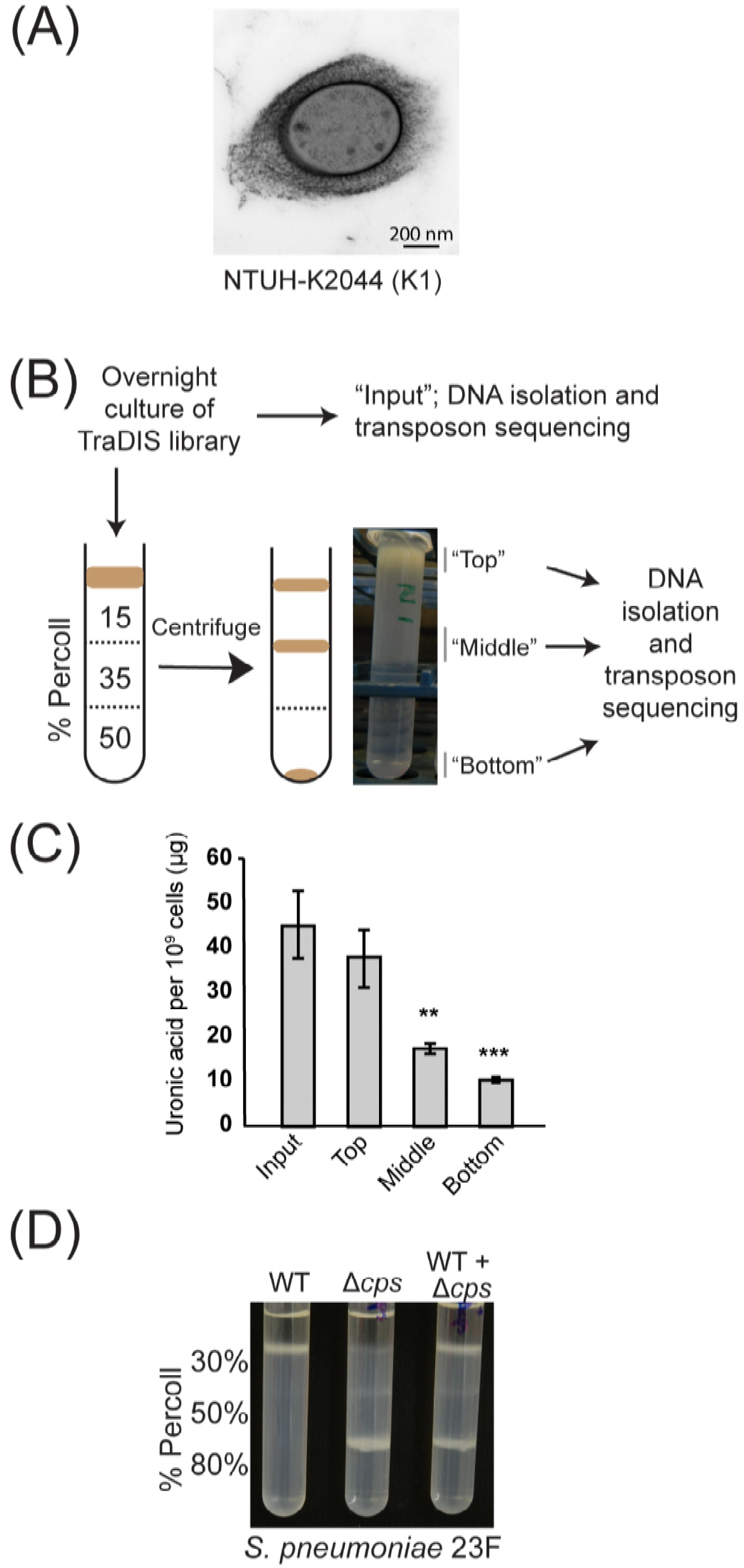
Summary of density-dependent TraDISort strategy. (A) Electron microscope image of capsulated *K. pneumoniae* NTUH-K2044. (B) Schematic of the TraDISort strategy to identify capsule regulators. A high-density transposon library is applied to the top of a discontinuous Percoll gradient, which is then centrifuged at moderate speed to separate capsulated and non-capsulated mutants. The separate fractions are sequenced to identify transposon-gDNA junctions. (C) Validation of the Percoll gradient method for separating cells by capsule phenotype. Individual fractions immediately following separation on a Percoll gradient were adjusted to OD_600_ = 4 in sterile PBS, and assayed for uronic acid content. Statistical significance was evaluated by one-way ANOVA followed by Tukey’s Honest Significant Differences (HSD) test, and is reported for each fraction relative to the input sample (** p < 0.01; *** p < 0.001). (D) Application of density separation method to *S. pneumoniae* strain 23F.

We then tested this method with a high-density transposon insertion library of *K. pneumoniae* NTUH-K2044 (generated as described in Materials and Methods). This mutant library gave rise to three fractions on a 15%-35%-50% Percoll gradient (Fig. 1B). The middle and bottom fractions separated from the NTUH-K2044 library contained less capsule than the input library culture, as shown by quantification of capsular uronic acids (Fig. 1C) and the hypermucoidy centrifugation test (Fig. S1C). To determine whether altered migration in a Percoll gradient was the result of stable mutant phenotypes, the fractions of the *K. pneumoniae* NTUH-K2044 gradient were cultured overnight and centrifuged on a fresh gradient. The “bottom” and “top” fractions migrated to the same position as before, while the “middle” fraction showed a partially heritable phenotype and was distributed across the top and middle positions (Fig. S1D).

These data indicated that centrifugation on a discontinuous density gradient successfully separated *K. pneumoniae* populations based on their capsule production, and that variations in density among library mutants were, in part, due to stable phenotypes. Random-prime PCR of several colonies from the bottom fraction of the *K. pneumoniae* NTUH-K2044 library identified insertions in known capsule biosynthetic or regulatory genes (Supplementary text S1), and a *wza* insertion mutant was retained as a capsule-negative control for future experiments. We then wished to determine whether our method could be used for capsule-based separations of other species, for which we tested capsulated and non-capsulated *Streptococcus pneumoniae* 23F. These strains were separated reproducibly on a Percoll gradient (Fig. 1D), indicating that our method is applicable to other bacterial species.

### Density-TraDISort identifies multiple capsule-associated genes in *Klebsiella pneumoniae*

TraDISort is a term for transposon sequencing screens based on the physical, rather than survival-based, enrichment of insertion mutants from saturated libraries (48), which allows examination of phenotypes that are not linked to survival. We call our approach density-TraDISort to distinguish it from the original method using fluorescence-based flow sorting as the physical selection method. The final density-TraDISort approach is shown (Fig. 1B). Briefly, transposon mutant libraries were grown overnight at 37 °C in LB, applied to the top of a Percoll gradient and centrifuged at a moderate speed for 30 minutes (see Materials and Methods). Following centrifugation, each bacteria-containing fraction was extracted and subject to TraDIS, as was a sample of the input culture. Data were analysed using the Bio-TraDIS pipeline (see Materials and Methods). The *K. pneumoniae* NTUH-K2044 library contained approximately 120,000 unique transposon insertion sites (equivalent to an insertion every 45 bp), with a median of 14 insertion sites per gene. Statistics summarising the sequencing results from each fraction are reported in Table S2.

Unlike traditional growth-based transposon-insertion screens, our density-TraDISort setup combines positive and negative selection within a single experiment: mutants with reduced capsule can be identified through their loss from the top fraction, or by virtue of their enrichment in another fraction (see Fig. 2A for an example). We applied stringent cutoffs based on both selections to identify putative capsule-related genes. Briefly, a gene was only counted as a hit if it was (i) lost from the top fraction (log_2_FC < -1, q-value < 0.001) and (ii) enriched in another fraction (log_2_FC > 1, q-value < 0.001) (see Tables S3, S4). The presence of a “middle” fraction also enabled us to identify genes which increased capsulation when disrupted (example in Fig. 2B). These were apparent in the raw TraDIS plot files as genes where all or nearly all the mutants localised to the top fraction following centrifugation (mutants with wild-type capsule were distributed between the top and middle fractions, in keeping with the migration pattern of wild-type NTUH-K2044 (Fig. S1A)). We defined putative “capsule up” genes as those which showed a marked depletion in the middle fraction relative to the input (log_2_FC < -3; q-value < 0.001), with genes with very low initial read counts (log_2_CPM < 4) excluded (Tables 1, S4). Examples of TraDISort data for several genes known to affect capsule production are shown in Figure S2C.

**Table 1.**
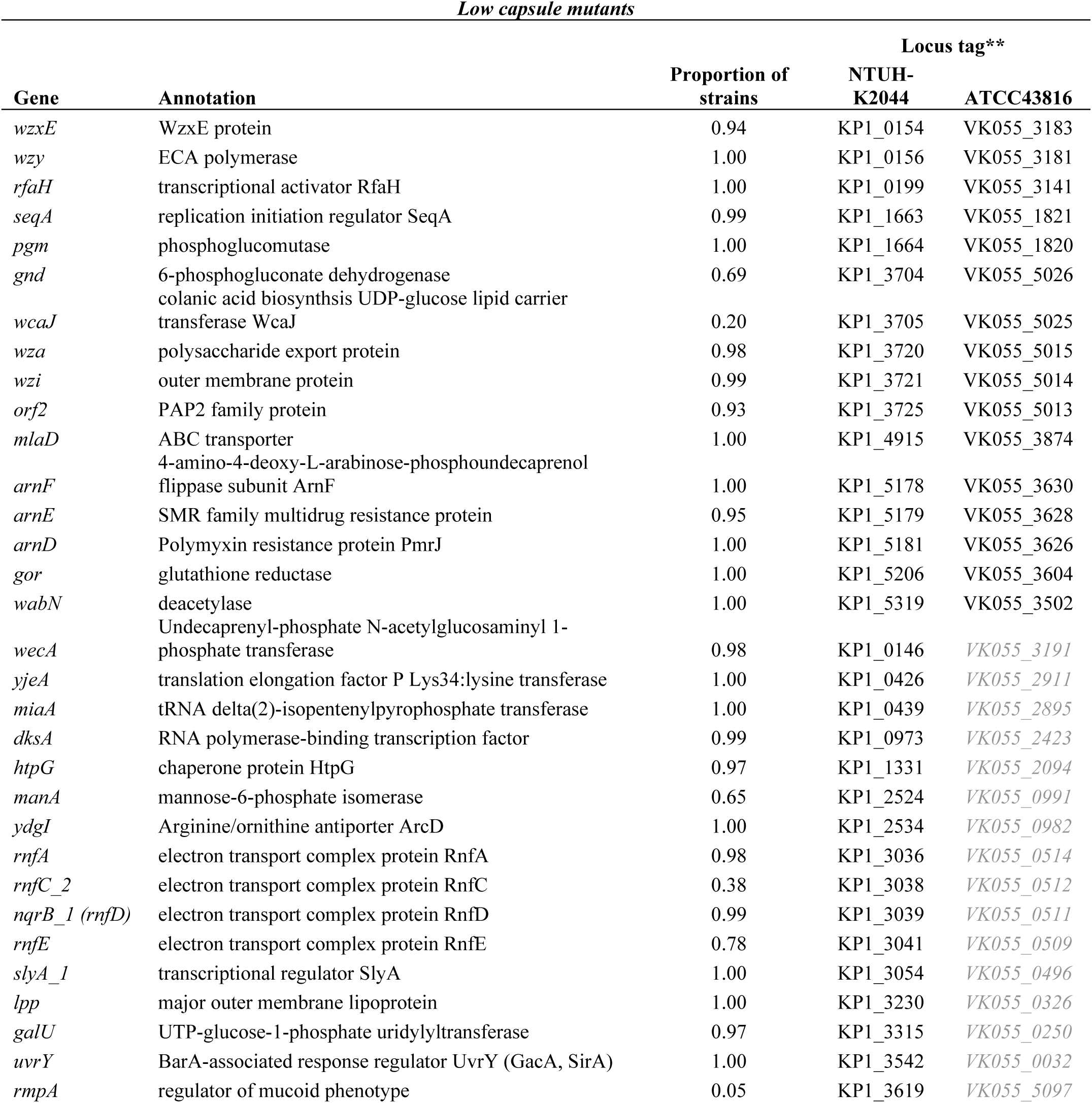
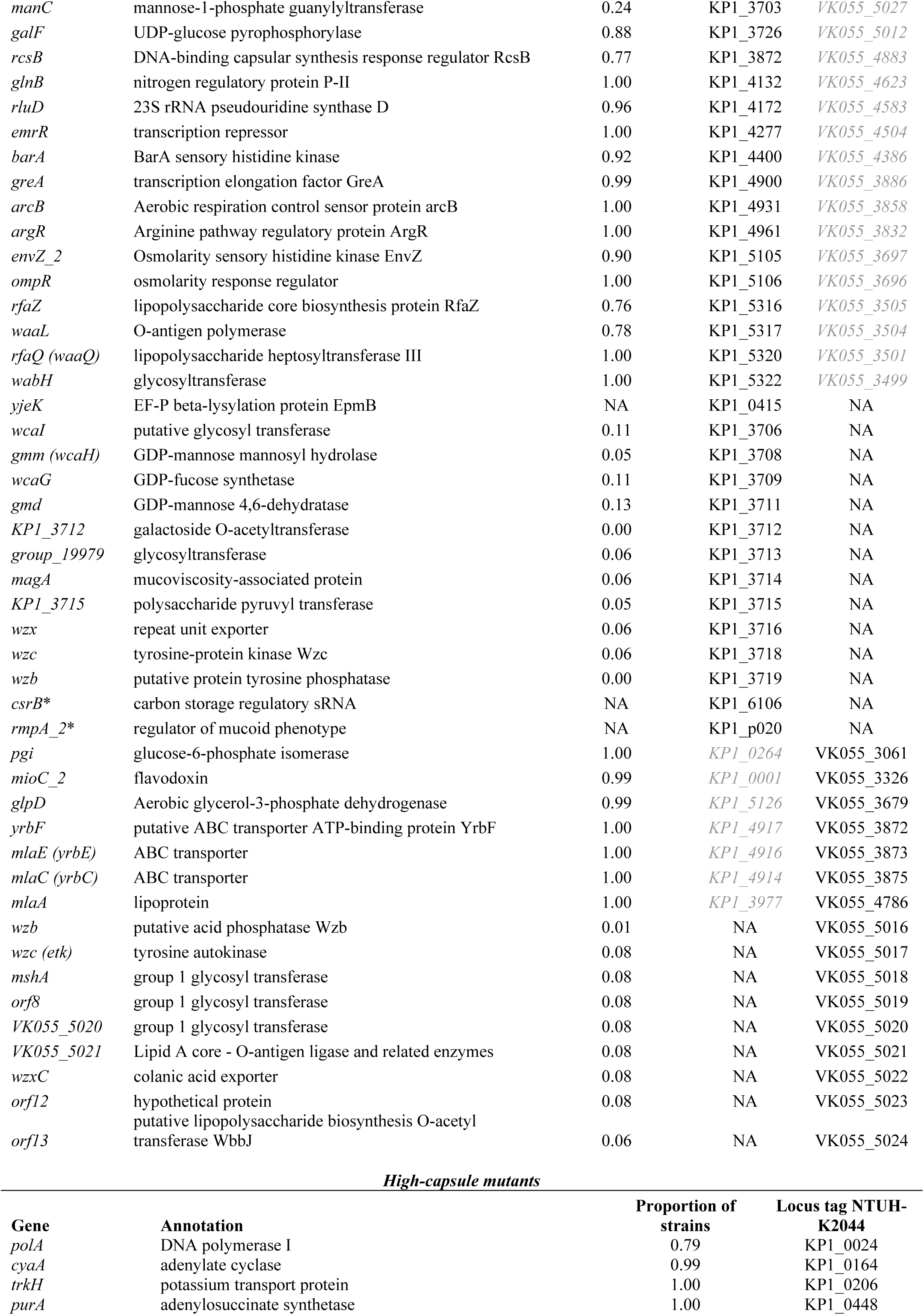
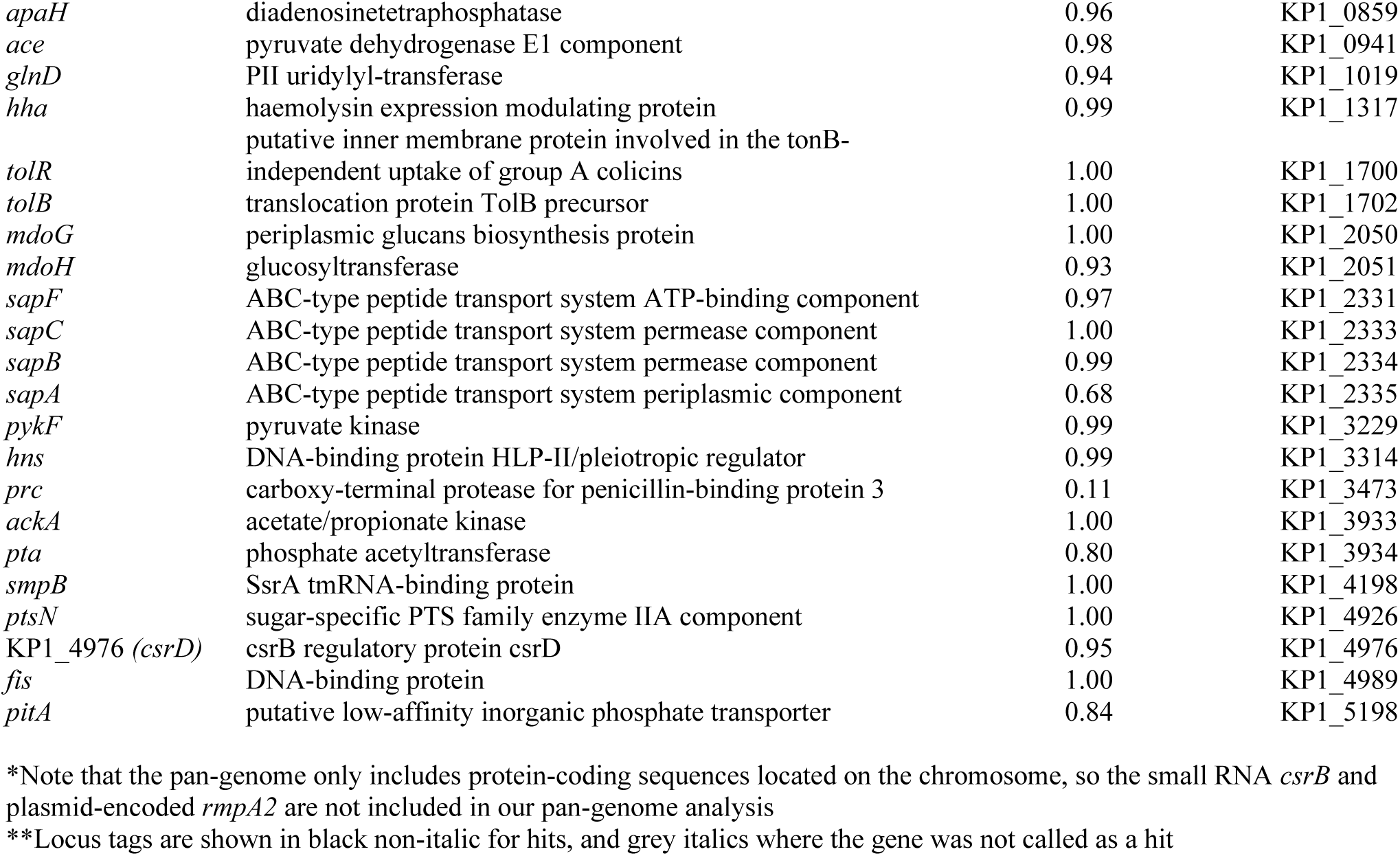
Genes in which mutants have altered capsule production. A list of all statistically-significant genes from this TraDIS screen which, when disrupted by transposon insertion, increase capsule production in *K. pneumoniae*. Cutoff criteria are described in Methods. Gene names and functional annotations are taken from the pan-genome consensus file (see Materials and Methods). The complete dataset, including statistical data, is provided in Tables S3 and S4.

**Figure 2:**
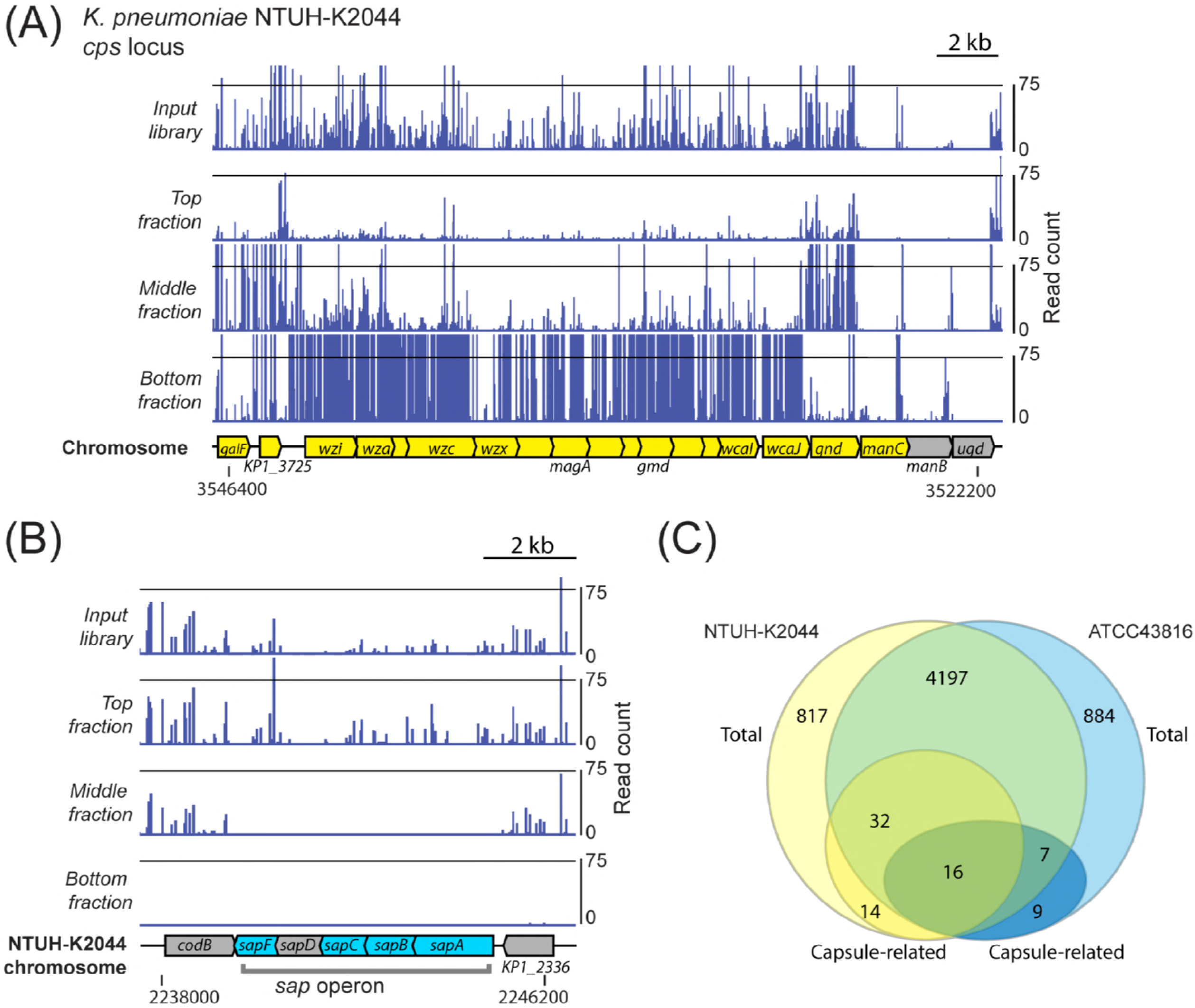
(A) Results of TraDIS mapping at the capsule locus of *K. pneumoniae* NTUH-K2044. Transposon insertions in capsule genes were very abundant in the input sample. These majority of these mutants were not found in the top fraction but were instead enriched in the middle fraction (e.g., *gnd, galF*) or the bottom fraction (all genes from *wcaJ* to *wzi*). Genes defined as hits in our screen are shown in yellow, others in grey. (B) Insertion mutants in genes of the *sap* ABC transporter locus were not found in the middle fraction, suggesting higher capsule production than wild-type. Genes which are putative increased capsule hits are shown in blue, with others in grey. Note that *sapD* had a very low number of reads in all fractions. The flanking genes, *codB* and KP1_2336, are examples of genes where transposon insertion does not change capsule, and the insertion mutants are found in both the top and the middle fractions. (C) Common and strain-specific genes required for full capsule production in *K. pneumoniae* NTUH-K2044 and ATCC43816. Putative increased capsule mutants are not considered in this chart. These two strains share approximately 4,200 genes, and the majority of genes required for full capsule production are present in both strains. Sixteen genes were defined as capsule-related in both strains. Note that the gene content of the *cps* locus differs between these two strains, which accounts for the majority of hits in strain-specific genes.

In total, we identified 62 genes required for full capsule production in *K. pneumoniae* NTUH-K2044 (Tables 1, S4). We also identified 26 putative “capsule up” mutants in this strain (Tables 1, S4). The biological roles of the genes determined to influence capsule production are discussed in more detail below.

### Global regulators, metabolic genes and cell surface components affect capsule in *K. pneumoniae*

Our “capsule down” mutants included almost every gene of the capsule biosynthesis locus of *K. pneumoniae* NTUH-K2044 (Fig. 2A), further confirming that our experimental strategy had successfully isolated capsule-deficient mutants. The *manB* and *ugd* genes were not called as hits, however these contained very few insertions in the input sample (visible in the plot files above these genes in Fig. 2A). In addition to biosynthetic genes, the known capsule regulators *rmpA, rcsB* and *rfaH* were identified (Fig. S3C).

Other “capsule down” hits included multiple cell surface components, metabolic genes and genes of global regulatory systems. Extended functional information for all of our hits, including reported links to capsule production in *Klebsiella* and other proteobacteria, is included in Table S4. Gene enrichment analysis using the TopGO package (49) indicated that genes for polysaccharide metabolic processes (p = 1.7 × 10^-5^), molecular transducers (p = 0.0042) and protein kinases (p = 0.005) were overrepresented in our hits (all probabilities are from Fisher’s exact test, parent-child algorithm (50)).

Several global regulators were identified as affecting capsule production. These included the anaerobic/redox-sensitive sensor kinase ArcB, the osmotic stress response system OmpR-EnvZ, the BarA-UvrY system, the arginine repressor ArgR, and the carbon storage regulatory small RNA CsrB. The transcription regulators MprA and SlyA were also needed for full capsule production. Metabolic genes were also identified, including genes of the electron transport chain, glycolytic enzymes, and the nitrogen source regulatory protein GlnB.

The “capsule up” mutants of NTUH-K2044 included nucleoid proteins, membrane-bound transporters such as SapBCDF (Fig. 2B), and several metabolic genes (Table S4). Interestingly, some of the “capsule up” hits are known antagonists or regulators of genes in the “capsule down” hits: CsrD (which targets CsrB for degradation), H-NS (reported to be antagonised by SlyA), and GlnD (modifies and controls activity of GlnB).

Finally, our results also included many genes for other cell surface components; genes for the synthesis and modification of LPS were identified, along with enterobacterial common antigen genes, and the lipoprotein Lpp.

### Distinct and overlapping capsule regulators in two *K. pneumoniae* strains

We wished to determine the extent of conservation of our capsule-regulatory hits across the *Klebsiella* species, and the overlap between the capsule-influencing genes in different *Klebsiella* strains. To explore this question, we applied our density-TraDISort method to a second strain, *K. pneumoniae* ATCC43816, a capsule type K2 strain which is commonly used in *Klebsiella* infection studies (Fig. S3A). We constructed a *K. pneumoniae* ATCC43816 saturated transposon insertion library with ∼250,000 unique insertion sites (or an insertion every 22 bp) and a median of 36 insertion sites per gene (see Materials and Methods; Fig. S3C-D). This library resolved into two fractions on a 35%-50% Percoll gradient, with no obvious heritable “middle” fraction, and fractions were collected and their transposon insertion junctions sequenced as for *K. pneumoniae* NTUH-K2044. Genes were counted as hits in this strain if they were lost from the top fraction relative to the input (log_2_FC < -1, q-value < 0.001) and enriched in the bottom fraction (log_2_FC > 1, q-value < 0.001). The lack of a “middle” fraction in this experiment means there is less sensitivity for identifying reduced-capsule mutants, particularly using the second, positive selection-based filtering criterion. We used a pan-genome generated from 263 annotated *K. pneumoniae* genomes (Table S5) to define the common genes in NTUH-K2044 and ATCC43816 and determine their prevalence across the *K. pneumoniae* population (Tables 1, S4).

We identified 34 candidate capsule-regulatory genes in *K. pneumoniae* ATCC43816. As observed for NTUH-K2044, the genes of the capsule biosynthetic locus were nearly all called as hits (Fig. S3E). Of the three genes that were not, two had very low initial insertion counts, and the third met our first selection criterion of being lost from the top fraction but was not enriched in the bottom fraction. Putative capsule-influencing genes of ATCC43816 fell into diverse functional categories, with cell surface components, metabolic genes, transporters and known regulators implicated.

Although the majority of capsule hits are encoded in the core genome (48 of 62 in *K. pneumoniae* NTUH-K2044; 23 of 32 in *K. pneumoniae* ATCC43816, Fig. 2C), only sixteen genes were called as hits in both strains. These shared genes included five shared components of the two strains’ capsule biosynthesis loci, the transcription antiterminator *rfaH*, two enterobacterial common antigen genes, three genes of the *arn* operon responsible for modification of LPS lipid A with L-Ara4N, and the glutathione reductase *gor*. Strain-specific differences were also identified, with the caveat that NTUH-K2044 produces more capsule (see Fig S3B) and therefore afforded us greater sensitivity in our experiment. Genes that were identified as required for ATCC43816 capsule production but not NTUH-K2044 included the *yrbCDEF* (*mlaCDEF*) ABC transporter and the *mlaA* gene, which are involved in maintaining outer membrane asymmetry through the cycling of phospholipids (51). Thirty-two genes common to both strains had a “capsule down” phenotype in NTUH-K2044 but were not called as hits in ATCC43816 (note that “capsule up” hits are not included in our comparison as these were not resolved in ATCC43816). These included the electron transport pump *rnf*, global regulators such as *arcB* and *ompR*, the transcription factor *mprA*, and several additional cell surface component biosynthetic genes. It appears that for at least some conserved genes their influence on capsule is strain-specific.

### Phenotypes of single-gene mutants confirm results of density-TraDISort

We generated a set of ten single-gene deletion mutants in *K. pneumoniae* NTUH-K2044, and three in *K. pneumoniae* ATCC43816, in order to validate the results of our density-TraDISort screen. Genes selected for mutagenesis were the known capsule and LPS regulator *rfaH* in both strains, the LPS O-antigen ligase *waaL*, the aerobic respiration control sensor *arcB*, and multiple transcriptional regulators (*ompR, argR, slyA, mprA, uvrY*). We also deleted the *arnF* gene in ATCC43816. The *sapBCDF* ABC transporter in NTUH-K2044 was examined in order to validate our assignment of “capsule up” hits.

Single gene knockout mutants were grown under the same conditions as in the original screen and subjected to density gradient centrifugation. Every mutant showed a banding pattern consistent with the results of density-TraDISort (Fig. 3A-B). The Δ*arcB* mutant appeared to have two populations, one with wild-type and one with reduced capsule, most other NTUH-K2044 mutants migrated to the position of the middle fraction, while the Δ*waaL*, i-*wza* and Δ*rfaH* mutants migrated to the bottom of the gradient. The Δ*sapBCDF* mutant showed higher density than wild-type and remained above the 15% Percoll layer, with no movement into the gradient itself. Mutants of NTUH-K2044 were also tested for hypermucoidy and uronic acid production, and these experiments showed increased capsule in NTUH-K2044 Δ*sapBCDF* and reduced capsule in all other mutants (Fig. 3C, Fig. S4B). Of the *K. pneumoniae* ATCC43816 mutants (Fig. 3B), Δ*rfaH* migrated to the bottom of the gradient and Δ*arnF* to the middle and bottom, while Δ*mprA* showed the same pattern as wild-type. This is consistent with the TraDISort assignment of this gene as having a strain-specific effect on capsule, at least in these growth conditions (Table 1).

**Figure 3:**
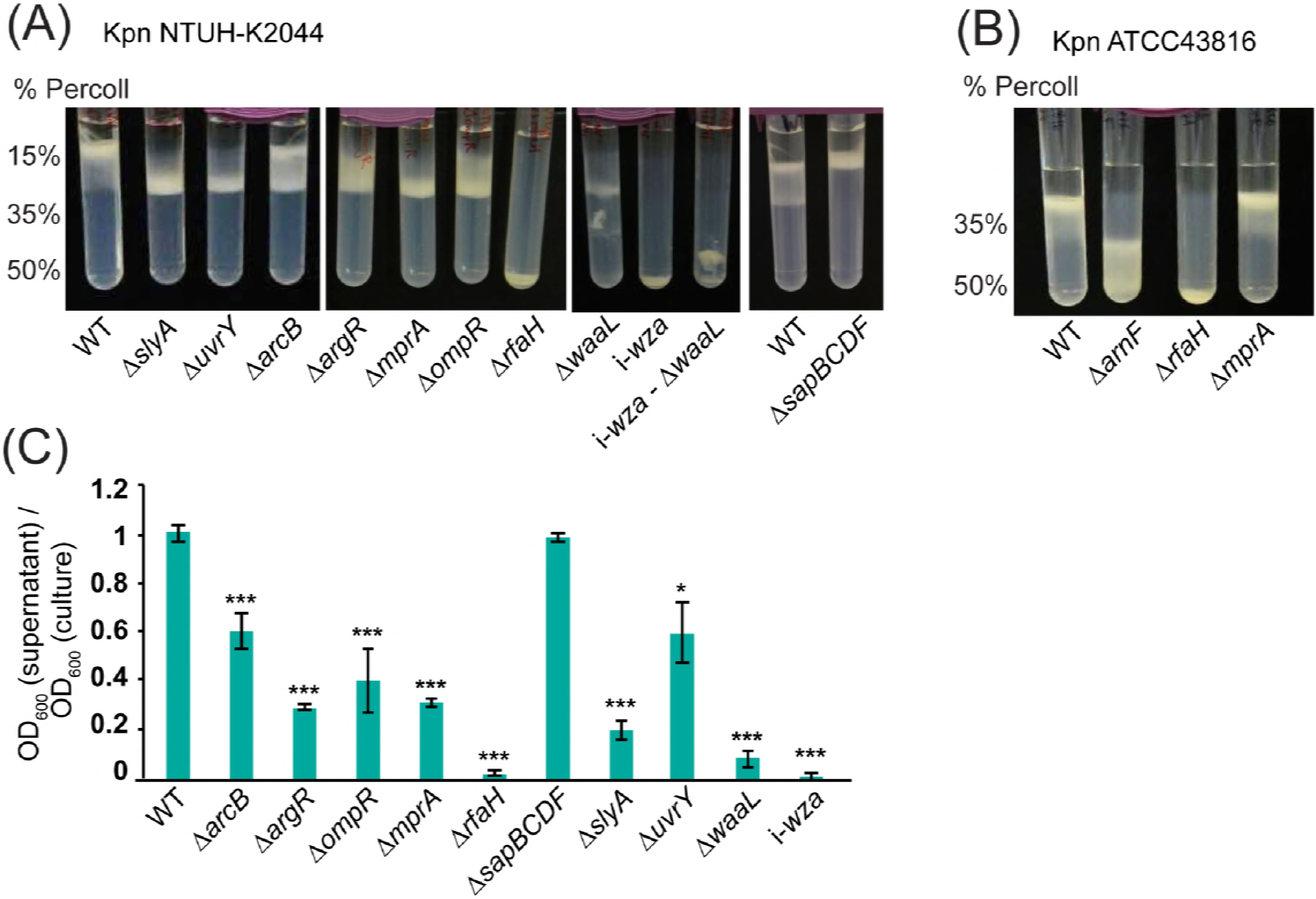
Validation of putative capsule regulators with single-gene deletion mutants. (A) Percoll gradient centrifugation of clean deletion mutants in selected NTUH-K2044 genes. All of the genes tested showed reduced density compared to wild-type, with the exception of the putative increased capsule mutant, Δ*sapBDEF*, which stayed above the 15% Percoll layer. (B) Validation of ATCC43816 deletion mutant phenotypes on 35%-50% Percoll gradients. The Δ*arnF* and Δ*rfaH* mutants showed reduced density compared to wild-type, while the Δ*mprA* mutant did not, in contrast to its phenotype in NTUH-K2044. (C) Hypermucoidy tests with Kpn NTUH-2044 mutants. Strains were grown to late stationary phase and cultures centrifuged for 5 min at 1,000 x *g*. The OD_600_ of the supernatant was measured and is presented here as a proportion of the starting OD_600_. *** p < 0.001, One-way ANOVA followed by Tukey’s HSD test, relative to wild-type.

Our hits included several genes with roles in LPS biosynthesis, which raised the possibility that LPS O-antigen may affect cell density independently of capsule. To test this, we constructed a Δ*waaL* deletion in our *wza* transposon insertion strain (see Materials and Methods). The resulting double mutant had no detectable reduction in density compared to the *wza* mutant on the standard 15%-35%-50% gradient or on 70% Percoll (the minimum concentration required to exclude the *wza* mutant, Fig. S4A), indicating that the LPS O-antigen alone does not affect the density of *K. pneumoniae*, at least within the resolution of this experiment.

### Virulence and capsule architecture of *K. pneumoniae* NTUH-K2044 ΔargR, ΔmprA, ΔsapBCDF and ΔslyA

We selected the transcription factors ArgR, SlyA and MprA, along with the ABC transporter SapBCDF, for further characterisation. ArgR represses arginine synthesis and transport as well as other genes (52), SlyA is an antagonist of H-NS (known to suppress capsule in *K. pneumoniae*) (53, 54), and MprA is a transcriptional regulator with an effect on capsule in UPEC (55). Both SlyA and MprA were very recently also shown to be virulence and capsule regulators in *K. pneumoniae*, and were renamed KvrA and KvrB (56). SapBCDF has been reported to mediate resistance to antimicrobial peptides (AMPs) in *H. influenzae* by importing them for degradation (57) and was presumed to have this activity in Enterobacteriaceae as well, though it has recently been reported that this pump functions as a putrescine exporter in *E. coli* and has no role in AMP resistance (58). ArgR and SapBCDF have not previously been linked to capsule regulation.

To confirm that the alterations in capsule production observed in the NTUH-K2044 Δ*argR*, Δ*slyA*, Δ*mprA* and Δ*sapBCDF* mutants were due to the deleted genes, each mutant was complemented by reintroducing the wild-type gene on the chromosome (see Materials and Methods). Complementation caused a complete restoration of wild-type capsule production, as measured by the hypermucoidy and uronic acid assays (Fig. 4A-B). To define changes in capsule architecture, each mutant strain was examined by transmission electron microscopy (Fig. 4C). Wild-type *K. pneumoniae* NTUH-K2044 had a thick, filamentous capsule of roughly half the cell diameter. The Δ*argR* and Δ*slyA* mutants had capsules with slightly reduced thickness and finer filaments, while the Δ*mprA* mutant had extremely fine and diffuse filaments such that the boundary of the capsule was not clear. The Δ*sapBCDF* capsule had some thick filaments but at lower density than NTUH-K2044, with an additional gel-like layer visible outside these filaments. The virulence of the Δ*argR*, Δ*slyA*, Δ*mprA* and Δ*sapBCDF* mutants, and their complements, was assessed by infection of research-grade *Galleria mellonella* larvae, an established invertebrate model for *Klebsiella* infections (59). Each of the reduced-capsule strains showed a virulence defect relative to the wild-type strain which was restored on complementation (Fig. 5A), while the Δ*sapBCDF* mutant did not have changed virulence compared to the wild-type.

**Figure 4:**
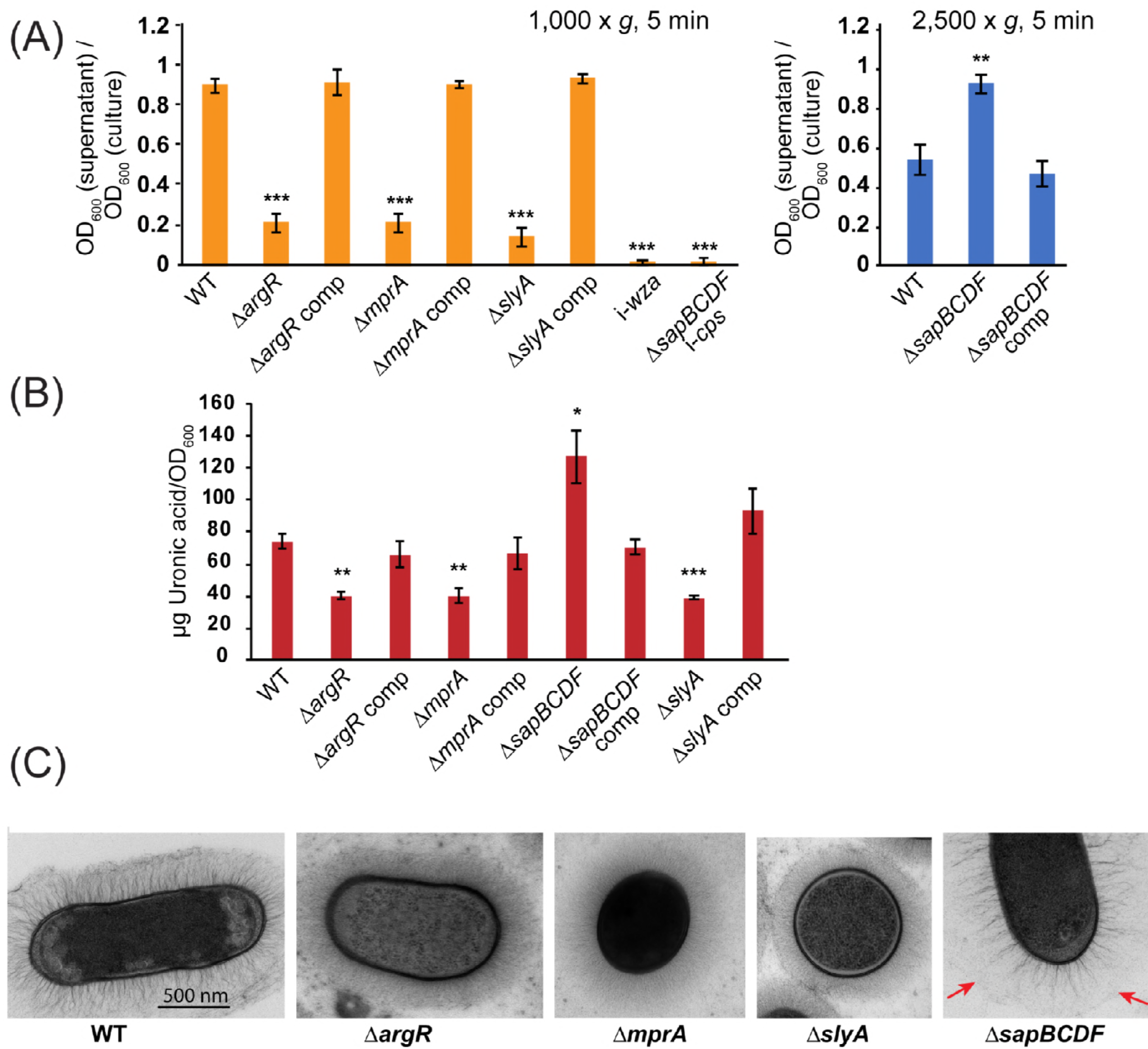
Complementation and electron microscopy of *K. pneumoniae* NTUH-K2044 Δ*argR*, Δ*mprA*, Δ*slyA* and Δ*sapBCDF* mutants. (A) Hypermucoidy assay. Strains were centrifuged at 1,000 x *g* for 5 min to define decreased hypermucoidy relative to wild type, or 2,500 x *g* to identify increases in hypermucoidy relative to wild type. Significant differences are indicated by ** p < 0.01 or *** p < 0.001, one-way ANOVA and Tukey’s HSD test. This shows an independent experiment to Figure 3C. (B) Uronic acid assay to confirm the capsule phenotype of each strain. Differences relative to wild type were evaluated by pairwise one-way ANOVA with Benjamini-Hochberg correction for multiple testing * p < 0.05, ** p < 0.001, *** p < 0.0001. (C) Transmission electron microscopy images of *K. pneumoniae* NTUH-K2044 and its Δ*argR*, Δ*mprA*, Δ*slyA* and Δ*sapBCDF* mutants.

**Figure 5:**
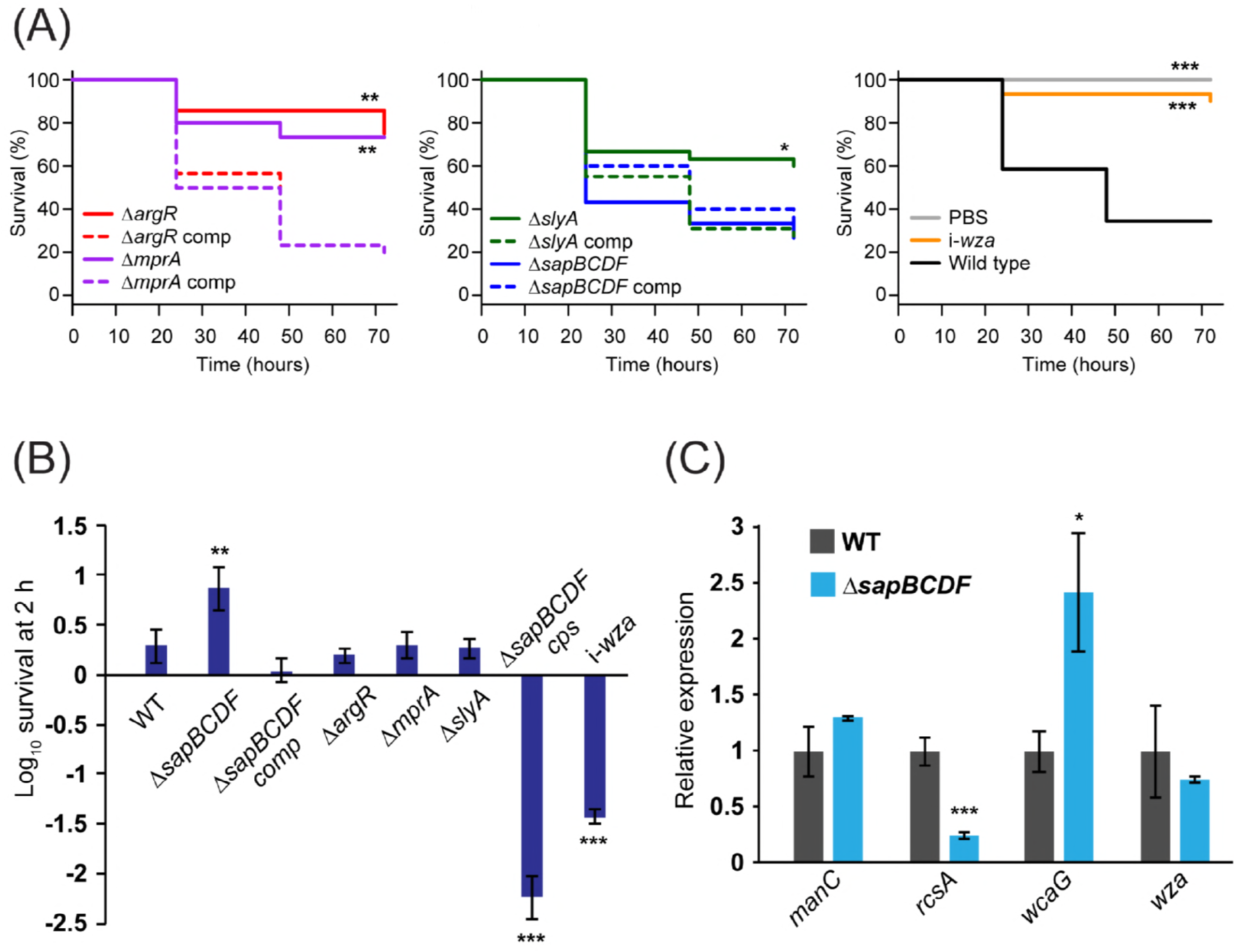
Virulence of selected mutants. (A) Killing of research-grade *Galleria mellonella* larvae by infection with *K. pneumoniae* NTUH-K2044 wild type or mutant strains. Larvae were infected at an inoculum of 10^5^. Differences in killing compared to wild type were evaluated using the Kaplan-Meier log rank test and are indicated by * p < 0.05, ** p < 0.01 or *** p < 0.001. (B) Survival in human serum. Differences relative to wild type are indicated by ** p < 0.01, *** p < 0.001, pairwise one-way ANOVA. (C) Expression of several capsule-related genes in NTUH-K2044 Δ*sapBCDF*. Transcript abundance was measured using the relative standard curve method with *recA* as a reference gene, and normalised to WT. * p < 0.05, *** p < 0.001, one-way ANOVA.

**Figure 6:**
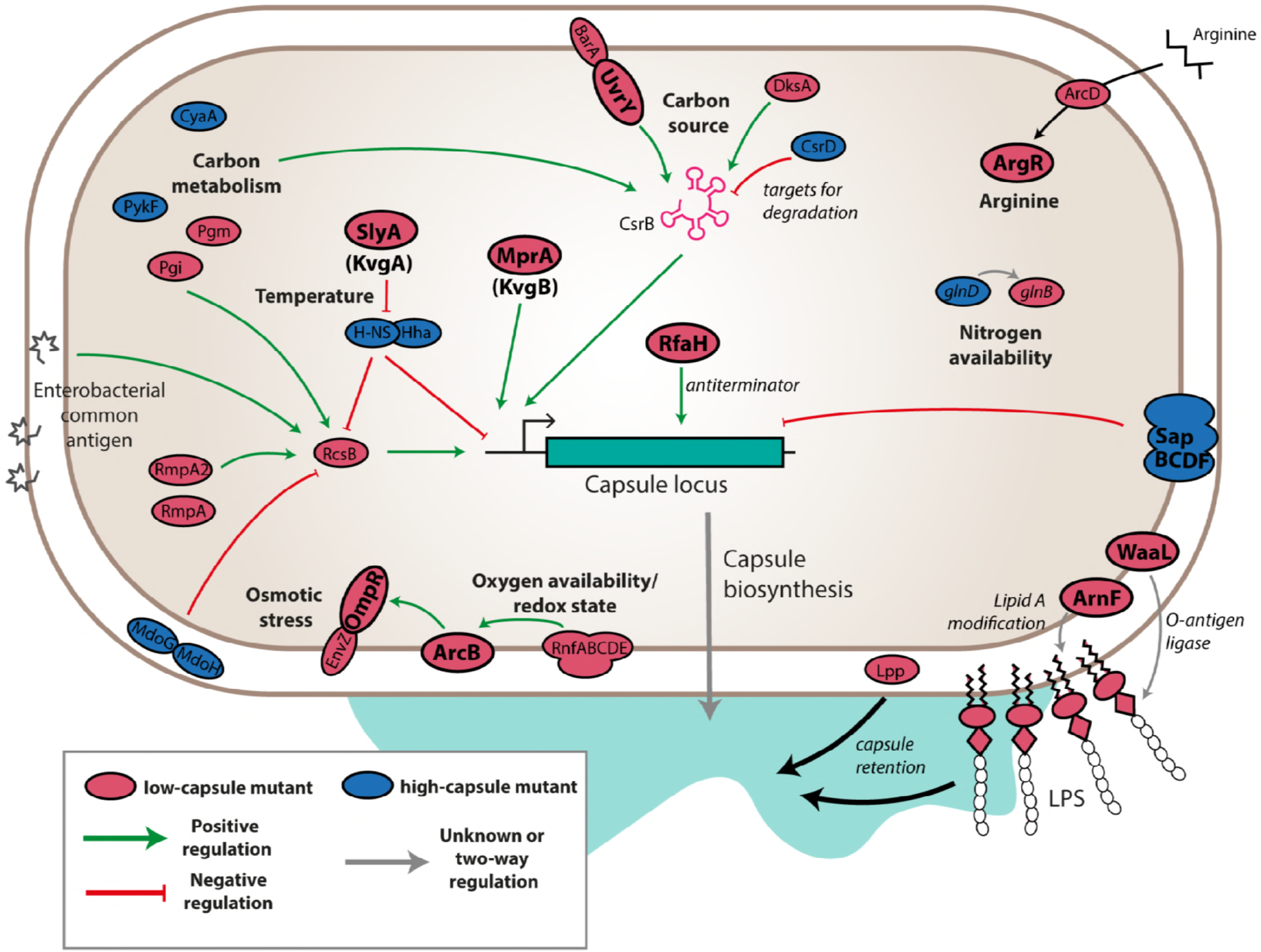
Overview of capsule regulation in NTUH-K2044. Products are coloured red for mutants with low capsule, blue for mutants with high capsule, and those genes that were validated in clean deletion knockouts are indicated with bold labels and outlines. Likely modes of action are indicated by green or red arrows for predicted positive and negative effects on transcription of the capsule locus. Grey arrows indicate inputs that may affect capsule synthesis without modulating transcription. Omitted are individual capsule biosynthetic genes, ECA biosynthetic genes, and components of the transcription and translation machinery.

### The Sap transporter alters serum survival but does not affect antimicrobial peptide resistance

We then examined the effect of ArgR, MprA, SlyA and SapBCDF on resistance to human serum. After two hours, NTUH-K2044 wild type showed full survival with a slight increase in viable count, while the *wza* mutant was reduced in viable count by ∼25-fold. The Δ*argR*, Δ*mprA* and Δ*slyA* mutants did not change significantly (Fig. 5B). The Δ*sapBCDF* showed increased viable count compared to the wild type strain, with a 7-fold increase over the course of the experiment. This increase was unexpected, as the wild type strain is already fully serum resistant. To determine if the increased serum survival was dependent on capsule production, a double Δ*sapBCDF cps* mutant was constructed (see Materials and Methods). This strain showed a drastic reduction in survival, suggesting that the NTUH-K2044 Δ*sapBCDF* mutant can grow better in serum because of its increased capsule (Fig. 5B).

We also tested the Δ*sapBCDF* mutant for resistance to the peptide antibiotics colistin and polymyxin B. The MIC was the same as that of the wild type, at 1 µg/ml for colistin, and 0.75 µg/ml for polymyxin B, indicating that the Sap transporter in *K. pneumoniae* does not contribute to antimicrobial peptide resistance.

### Sap mutation increases transcription of capsule middle genes without activating the Rcs system

We then wished to determine whether the mutation of the Sap transporter increased capsule production by acting on transcription. RNA was extracted from late exponential-phase wild type and Δ*sapBCDF* cells, and the abundance of three capsule locus transcripts – *manC, wcaG* and *wza* – was measured by qRT-PCR. These genes are transcribed from separate promoters. The Δ*sapBCDF* mutant showed elevated expression of *wcaG*, at 2.5 times wild type levels, while expression of *wza* and *manC* was not significantly changed (Fig. 5C). The Rcs phosphorelay system regulates capsule expression in *E. coli* and other enterobacteriaciae, and is induced by cues such as membrane stress (60, 61). RcsA is a component of the system which autoregulates and increases its own expression when activated. We measured *rcsA* transcript levels to determine whether loss of the Δ*sapBCDF* genes induces *rcsA* (Fig. 5C). Unexpectedly, levels of *rcsA* were much lower in the mutant than the wild-type, indicating that Sap-dependent induction of capsule expression does not occur through *rcsA*. Note, however, that RcsA is not required for all permutations of Rcs signalling as RcsB can interact with a number of partner proteins to regulate transcription (60).

### Capsule is at the centre of a complex regulatory network in *Klebsiella pneumoniae*

We identified numerous putative capsule regulators by density-TraDISort, and validated the results of our screen with single-gene deletion mutants. We propose an integrated model for how the genes we identified may collectively control *K. pneumoniae* NTUH-K2044 capsule. This model is based on our results and previous published work in *K. pneumoniae* and other Enterobacteria (particularly *Escherichia coli*). Full details of the literature relevant to each hit are in Table S4.

Major nodes for transcriptional control are the CsrB carbon source utilisation system and the Rcs phosphorelay system. Each of these systems is itself regulated by multiple genes identified in our study – CsrB integrates signals from the UvrY-BarA two component system (a “capsule down” hit) and various carbon metabolic genes, is activated by DksA, and is targeted for degradation by CsrD (a “capsule up” hit); the Rcs system is induced by MdoGH mutation, can cooperate with RmpA and RmpA2 to induce capsule, and also responds to carbon metabolism and some forms of enterobacterial common antigen. SlyA/KvrA and MprA/KvrB both promote capsule transcription (56). The SlyA/KvrA protein acts as a temperature-dependent switch which acts by relieving H-NS mediated transcriptional silencing; H-NS has been reported to operate on both the Rcs system and the capsule operon in *E. coli*, though the precise target of its activity in *Klebsiella* is not known. Capsule is also affected by the composition of the cell envelope, and mutations in *lpp* or various LPS-related genes can reduce the retention of capsule at the cell surface. Note that several genes related to cell envelope composition and membrane stress have been shown to regulate the Rcs system (61), therefore some of the cell envelope component genes identified in our study may act through RcsB. Our work has also uncovered novel regulators of capsule that, at this stage, cannot be tied to the wider regulatory network, such as *argR*, and the ABC transporter Sap. We intend to define the mechanisms by which these genes affect capsule in future studies.

## Discussion

We have developed a simple, robust technology for genome-wide studies of bacterial capsule, density-TraDISort, and applied it to identify capsule regulators in two strains of *K. pneumoniae*. In doing so, we have identified multiple positive and negative regulators of capsule production, including several genes not previously linked to capsule in this species. To our knowledge, this is one of the first studies employing a physical selection independent of bacterial survival and growth to separate TraDIS libraries, and is the first time that density-based physical selection has been applied to studying capsule regulation in *K. pneumoniae*. TraDISort/FAST-INSeq technology with fluorescence-based sorting has to date been used to identify genes affecting efflux of ethidium bromide, and mutations influencing expression of a *Salmonella* Typhi toxin reporter (48, 62). We have expanded the utility of this method by adding a selection step based on cell density, allowing us to resolve different capsulation states. We envisage that, in addition to facilitating genome-wide screens for altered capsulation in other bacterial species, density-TraDISort could be used to identify genes affecting cell size and shape, or aggregation.

Our study is the second application of TraDIS to screen for genes affecting bacterial capsule production, following a recent study focused on UPEC (55). The UPEC study utilised a capsule-specific phage to positively select transposon insertion mutants lacking capsule; two novel capsule regulators were identified in this way. Compared with phage-based selection, our method offers increased sensitivity – mutants with a range of capsule phenotypes can be identified, in addition to capsule-null mutations. In addition, there is the option for very stringent selection of hits, as cutoffs can be applied based on both negative selection (loss from the top fraction) and positive selection (enrichment in another fraction). However, density-based selection is less specific to capsule than phage infection, and there is the possibility that mutations could affect cell density in a capsule-independent manner. Interestingly, one of the novel *Klebsiella* capsule regulators identified in this study, MprA, was also shown to regulate capsule in UPEC. In *Klebsiella pneumoniae* this gene increases capsule production above a baseline in hypermucoid strains (Fig 3A, S4B) and (56)), while in UPEC an Δ*mprA* mutant did not produce capsule at all.

We have shown that capsule production in *K. pneumoniae* NTUH-K2044 is controlled by many different global regulatory systems, allowing us to describe a detailed snapshot of the control of capsule in this strain (Fig. 5). Note that our assay was performed on bacterial cells at late stationary phase growth, in LB medium, under microaerophilic conditions. This condition was used in this study because capsule production is high, offering good resolution for capsule-based selection of mutants. Additional regulators, linked to different cues and stresses, are likely to be involved in different environments. Many of the regulators identified in this study were only called as hits in the hypermucoid strain, *K. pneumoniae* NTUH-K2044. It remains to be seen whether these same regulators control capsule (though to a degree outside the resolution of our gradient) in other *K. pneumoniae* strains, though note that several genes of *K. pneumoniae* ATCC43816 (including *uvrY, barA, csrB, rcsA* and *rcsB*) met our first screening criterion of being lost from the top fraction but not the second of being enriched in the bottom fraction. We speculate that capsule production is subject to complex environmental control across the *Klebsiella* species, but that the hypermucoid phenotype is more expensive to maintain, and more sensitive to disruptions in its regulatory network.

Many of our hits are involved in the synthesis of other cell surface polysaccharides; these included genes for enterobacterial common antigen (ECA), as well as genes for the synthesis or modification of LPS. ECA is a non-immunogenic surface glycolipid found in various forms in Enterobacteriaceae, and structural modifications in this moiety can induce the Rcs system (63, 64). LPS is a major contributor to *K. pneumoniae* pathogenesis in sepsis, though to a lesser extent in pneumonia (12, 18), and various LPS modifications have roles in immune modulation during infection (65, 66). We are confident that the LPS mutations identified in our study affect capsule retention or biosynthesis, rather than density *per se*, because 1) deletion of the O-antigen ligase *waaL* did not reduce cell density in an acapsular *K. pneumoniae* NTUH-K2044 strain (Fig. 3A); 2) some LPS biosynthesis genes, but not all, were hits in our screen; and 3) the glucuronic acid moieties on the core LPS polysaccharide are required for capsule retention in *K. pneumoniae* (67, 68). In both strains we studied, disrupting genes of the *arn* operon reduced capsulation (Table 1, S4). The *arn* operon is responsible for modifying lipid A of LPS with 4-amino-4-deoxy-L-arabinose to mediate resistance to peptide antibiotics (69), but has not previously been linked to capsule. The *arnEF* genes encode a flippase thought to translocate the modified arabinose across the cell membrane (70), while *arnD* is involved in its biosynthesis (71). We hypothesise that the reduced capsule of mutants of *arnD* and *arnEF* on capsule is independent of Lipid A modification, because other genes in this operon did not affect capsule, and because a previous study showed that Lipid A modification with L-Ara4N does not occur in cells grown in LB (66). Overall, our results hint at a high degree of interdependence between the three major surface polysaccharides of *K. pneumoniae*.

The *sap* ABC transporter was found to promote capsule production when mutated, by increasing the expression of capsule middle genes (Fig. 3A, Fig. 4A, Tables 1, S3, S4). To our knowledge our study is the first to implicate *sapABCDF* in capsule regulation, though its full functions, or indeed the substrate of this transporter in *Klebsiella*, are not known. The *H. influenzae* Sap homologue mediates resistance to antimicrobial peptides by importing them for degradation, and is also required for haem uptake (57, 72), while in *E. coli* the Sap pump is reported to export putrescine and facilitate potassium import through TrkGH (58, 73). We found that the Sap transporter did not affect antimicrobial peptide resistance, which was also observed in *E. coli*. It is unclear how Sap mutation induces *wcaG* while suppressing an important component of the Rcs system – more work will be needed to define the role and mechanism of this transporter in *K. pneumoniae*.

We have developed a simple, broadly-applicable method for studies of capsulation, and used it to define the regulatory network that controls capsule in *K. pneumoniae* NTUH-K2044. We have also identified genes required for full production of capsule in a K2 strain. Although the majority of regulators are located in the core genome of *K. pneumoniae*, there are differences in the specific regulators deployed in the two strains we investigated, and it would be interesting to determine whether this pattern of strain-specific regulatory networks comprising primarily core genes holds across the *Klebsiella* phylogeny. This intra-species comparison, together with our data showing that density-based capsule selection can be used in other capsulated bacteria, also opens the possibility for robust interspecies comparisons of capsule regulation.

## Materials and Methods

### Culture conditions and microscopy

*K. pneumoniae* strains were cultured routinely in LB media supplemented with 1.5% w/v agar as appropriate. Cultures were supplemented with 12.5 μg/ml chloramphenicol and 12.5 μg/ml tetracycline when required. *S. pneumoniae* strains were grown on blood agar plates (Oxoid, CM02718) in microaerobic candle jars containing Campygen sachets at 37 °C, or in static BHI liquid media (Oxoid, SR0050C). The list of strains, plasmids, and oligonucleotides used in this study is reported in Table S1.

### Generation of transposon insertion libraries

TraDIS libraries were generated using the mini-Tn*5* transposon delivery plasmid pDS1028 (74), introduced into the recipient strain by conjugation. Full details are described in Supplementary methods.

### Mutant library fractionation on Percoll gradients

Bacterial mutant libraries were separated on the basis of their capsule expression by centrifugation on a discontinuous Percoll (GE Healthcare) density gradient for 30 minutes at 3,000 x *g* (Fig. 1B). Full details are described in Supplementary methods.

### Identification of transposon insertion sites by random-prime PCR

Genomic DNA was prepared from overnight cultures of single reduced-capsule mutants using the DNeasy® Blood and Tissue kit (Qiagen). Random-prime PCR to identify the transposon insertion site in each gDNA template was performed as described (75) using primers FS57-59 and FS109 and Herculase II polymerase (Agilent). Amplicons were sequenced using primer FS107.

### DNA extraction and next-generation sequencing

Genomic DNA (gDNA) was prepared from each Percoll-resolved fraction by phenol-chloroform extraction. From each gDNA preparation, 2 μg DNA was used to prepare TraDIS transposon-specific sequencing libraries as described previously, using primer FS108 for specific amplification of transposon junctions (43). Sequencing was carried out on the Illumina MiSeq platform using primer FS107.

### Analysis of TraDIS data

The analysis of TraDIS sequencing results was carried out using the Bio-TraDIS pipeline as described previously (43, 44), with minor modifications (see Supplementary methods). All scripts used in this study are available at https://github/sanger-pathogens/Bio-Tradis and https://github/francesca-short/tradis_scripts. Comparisons between fractions were based on normalised read counts per gene. Genes with (i) reduced mutant abundance in the top fraction, and (ii) increased mutant abundance in the middle or bottom fraction were called as decreased capsule hits, with thresholds of an absolute change in log_2_FC > 1 and q-value < 0.001. Increased capsule hits in NTUH-K2044 were defined as those that with severely reduced mutant abundance in the middle fraction (log_2_FC < -3; q-value < 0.001) without enrichment in the bottom fraction (log_2_FC < 1), with genes containing very few reads in any fraction excluded (*i.e.*, log_2_(counts per million) in the top fraction was greater than 4). Generation of the pan-genome, and enrichment analysis, is described in Supplementary methods.

### Construction of single-gene deletion strains

Single gene knockout mutants were constructed in *K. pneumoniae* by allelic exchange. Upstream and downstream sequences (> 500bp) for each target gene were amplified and joined by overlap PCR, cloned into pKNG101-Tc, and introduced into the recipient strain by conjugation with the *E. coli* β2163 donor strain. All primers used, and the resulting constructs, are listed in Table S1. Conjugation patches were incubated for one hour at 37 °C, then for 16 hours at 20 °C. Single-crossover mutants were selected on LB agar + 15 μg/ml tetracycline. Double-crossover mutants were selected on low-salt LB agar + 5% sucrose at room temperature, and were subsequently patched onto LB + sucrose and LB + tetracycline plates to confirm loss of the vector. Mutants were confirmed by PCR across the deleted region. Mutants were complemented by introduction of the relevant gene back into its original location on the chromosome by allelic exchange as described above, using a vector encoding the gene and its flanking region. The Δ*sapBCDF cps (KP1_3713)* double mutant was generated by random transposon mutagenesis of the Δ*sapBCDF* strain with the pDS1028 vector, followed by selection of acapsular mutants from the pool by density-gradient centrifugation and random-prime PCR to identify the insertion site.

### Quantification of capsule by uronic acid assay

Capsule extraction and quantification of uronic acids was performed as described previously (14, 76), with modifications (see Supplementary methods).

### Hypermucoviscosity assay

Cultures of *K. pneumoniae* were grown overnight in 5 ml LB medium at 37 °C. These cultures were sedimented at 1,000 x *g* or 2,500 x *g* for 5 minutes (room temperature). The OD_600_ of the top 500 μl of supernatant was determined by spectrophotometry. Results were expressed as a ratio of the supernatant OD_600_ relative to the input culture.

### Electron microscopy

Colonies were taken directly from an agar plate, high-pressure frozen in a Balzers HP010, and freeze-substituted for 8 hours in acetone containing 0.1% tannic acid and 0.5% glutaraldehyde at -90 °C followed by 1% osmium tetroxide in acetone for 24 hours at -50°C. They were then embedded in Lowicryl HM20 monostep resin. Ultrathin sections were cut on a Leica UC6 ultramicrotome and contrasted with uranyl acetate and lead citrate. Images of bacteria were taken on an FEI Spirit Biotwin 120 kV TEM with a Tietz F4.15 CCD.

### Serum resistance assay

Bacteria were grown in LB to an OD_600_ of 1, pelleted and resuspended in sterile PBS. Human sera (400 µl; Sigma-Aldrich S7023) was pre-warmed to 37 °C and added to 200 µl bacterial suspension, and the mixture was incubated at 37 °C for two hours. Viable bacterial counts were determined before and after incubation.

Galleria mellonella *infection*

Larvae of *G. mellonella* were purchased from BioSystems Technology UK (research-grade larvae) and used within one week. Bacteria were grown overnight, subcultured and grown to an OD_600_ of 1, then resuspended in sterile PBS. Larvae were infected by injecting the bacterial suspension (10^5^ cells) into the right hind proleg of the larvae using a Hamilton syringe. Infected larvae were incubated at 37 °C and monitored every 24 h, and were scored as dead when they were unresponsive to touch. Thirty larvae were used per strain, and these were infected in three batches of 10 using replicate cultures.

### Antimicrobial peptide resistance tests

Strains were grown overnight, subcultured and grown to an OD_600_ of 1.0. This culture (100 µl) was spread on the surface of an LB agar plate, dried, and an E-test (BioMerieux) strip was placed on the surface of the plate. Plates were incubated face-up at 37 °C and the result was read after 6 hours to avoid overgrowth of the capsule interfering with the reading.

### RNA extraction and qRT-PCR

Bacteria were grown in LB medium at 37 °C to an OD_600_ of 1.0. Cultures were mixed with 2 x volumes RNAProtect reagent, centrifuged, and RNA extracted using the Masterpure Complete DNA and RNA Purification kit (Epicentre) according to the manufacturer’s instructions. Samples were then subject to in-solution DNaseI digest (Qiagen), and cleaned up using the Qiagen RNeasy Mini kit. Reverse transcription of 200 ng RNA was performed using the ProtoScriptII enzyme (NEB) with supplied instructions. Transcripts were quantified using a StepOne real-time PCR instrument with the KAPA SYBR FAST qPCR kit. Relative abundance was determined using the relative standard curve method with *K. pneumoniae* NTUH-K2044 gDNA as a standard, and *recA* as the reference gene (77).

## Accession numbers

Sequences generated during this study have been deposited into the European Nucleotide Archive (ENA; http://www.ebi.ac.uk/ena) under study accession number ERP105653.

## Acknowledgements

We thank Matt Mayho, Jacqui Brown, and the sequencing teams at the Wellcome Trust Sanger Institute for TraDIS sequencing and the Pathogen Informatics team for advice on the analysis. We also thank Susannah Salter, Nick Croucher, George Salmond, Rita Monson, Jose Bengoechea and Jin-Town Wang for supplying strains and plasmids. This work was supported by Wellcome (grant 206194). M.J.D. is supported by a Wellcome Trust Sanger Institute PhD Studentship. F.L.S. is supported by a Sir Henry Wellcome postdoctoral fellowship (grant 106063/A/14/Z).

## Supplementary material legends

**Figure S1.** Validation of density gradient centrifugation for capsule-based separations. (A) *K. pneumoniae* NTUH-K2044 (K1), *K. pneumoniae* ATCC43816 (K2) and *E. coli* DH5α grown at 37 °C each localised to a different location on a 15%-35%-50% Percoll gradient, and the two *Klebsiella* strains migrated further down the gradient when grown at 25 °C. Locations equivalent to the Top, Middle and Bottom fractions used in (B) are indicated by red letters ‘T’, ‘M’ and ‘B’. A non-hypermucoid capsule type K106 strain, *K. pneumoniae* RH201207, was distributed throughout the 35% Percoll layer following centrifugation. (B) Viable counts of gradient fractions of wild type *K. pneumoniae* NTUH-K2044 and *K. pneumoniae* ATCC43816. The NTUH-K2044 strain localised primarily above the 15% Percoll layer, with a smaller proportion of cells in the middle fraction (4% of the viable count of the top fraction). The viable count of the bottom fraction was close to the detection limit (indicated by a dashed line) at 0.0005% of that of the top fraction. The ATCC43816 strain migrated almost solely to the 15%-35% interface of the gradient, with 0.01% of cells located in the bottom fraction in this experiment. *** p < 0.001, one-way ANOVA followed by Tukey’s HSD performed on log-transformed viable counts. (C) Hypermucoidy test of individual fractions of the NTUH-K2044 TraDIS library following separation. (D) Heritability of cell density phenotype following overnight growth of fractions. The “middle” fraction had a partially heritable phenotype and was distributed across the previous locations of the “top” and “middle” fractions. The top and bottom fractions localised to the same place following overnight growth.

**Figure S2.** (A) Relationship between sequencing depth and number of insertion sites in the *Klebsiella pneumoniae* NTUH-K2044 library, sequenced input sample 1. The plot was generated using the **seq_saturation_test.py** script available at https://github.com/francesca-short/tradis_scripts. (B) Gene-wise insertion index values along the chromosome of *K. pneumoniae* NTUH-K2044. (C) Density-TraDISort results for NTUH-K2044 genes with known capsule phenotypes. Mutants with unchanged capsule production were located in the top and middle fractions, as shown for *fadB*. Severe capsule reduction resulted in migration to the bottom fraction, shown for *rfaH*, while mutants in genes required for the hypermucoidy phenotype (*rcsB* and *rmpA*) were located primarily in the middle fraction. Genes where mutation results in increased capsule were evident as those that were absent from the middle fraction, but enriched in the top fraction, as shown for *hns*.

**Figure S3.** TraDIS analysis of capsule regulation in *Klebsiella pneumoniae* ATCC43816 (A) Electron microscope image of ATCC43816. (B) Uronic acid assay to validate density-based separation for this strain and compare its capsule production to NTUH-K2044. ** p < 0.01, one-way ANOVA and Tukey’s HSD. (C) Relationship between sequencing depth and number of unique insertion sites identified in the *Klebsiella pneumoniae* ATCC43816 mutant library. (D) Distribution of insertion sites across the chromosome. (E) TraDIS plot files at the capsule locus of *K. pneumoniae* ATCC43816; almost all mutants were found in the bottom fraction. Genes called as capsule-regulatory hits are shown in yellow, those not called as hits are in grey.

**Figure S4.** (A) Mutation of *waaL* does not further reduce density in a *wza* mutant strain. The indicated mutants of NTUH-K2044 were centrifuged on a 70% Percoll layer, which was determined to be the concentration required to retain the *wza* mutant. (B) Uronic acid assay with mutants of NTUH-K2044. These data are from the same experiment as that shown in Figure 4B. Differences relative to wild type were evaluated by pairwise one-way ANOVA with Benjamini-Hochberg correction for multiple testing (* p < 0.05, ** p < 0.01 or *** p < 0.001).

**Supplementary Text S1.** Extended methods, supplementary references, and random-prime PCR identification of transposon insertion sites in acapsular clones.

**Supplementary Table S1**. Strains, plasmids, and oligonucleotides used in this study.

**Supplementary Table S2.** Summary statistics and reference numbers for TraDIS samples sequenced in this study.

**Supplementary Table S3.** Raw comparison data for *K. pneumoniae* ATCC43816 and NTUH-K2044 capsule gradient fractions.

**Supplementary Table S4.** Combined capsule hits in *K. pneumoniae* ATCC43816 and NTUH-K2044 with pan-genome information and references to relevant literature.

**Supplementary Table S5.** List of genomes used to generate the *K. pneumoniae* pan-genome.

